# “Loss of circulating glucocorticoid rhythm disrupts the circadian transcriptome and vascular reactivity in the mouse renal artery.”

**DOI:** 10.1101/2025.04.29.650449

**Authors:** Georgios Krilis, Hannah M. Costello, Frances Turner, Robert Little, Anastasiia Komolova, Josselin Nespoux, Alicja Czopek, Donald R. Dunbar, Matthew A. Bailey, Jess R. Ivy

## Abstract

**Aims:** Molecular clocks in the vasculature contribute to the normal circadian blood pressure (BP) rhythm. Glucocorticoids are known to synchronise molecular clocks within peripheral tissues. Disruption of the endogenous glucocorticoid rhythm causes “non-dipping”, a BP pattern associated with elevated cardiovascular risk. The mechanisms for this are unclear but vascular changes likely contribute. We examined the effect of impaired glucocorticoid rhythmicity on the time-of-day dependent vascular function and define the circadian transcriptome under control conditions and under arrhythmic glucocorticoids.

**Methods and Results:** Male C57BL6J mice kept on a 12:12 hour light/dark cycle were implanted with a subcutaneous slow-release pellet containing vehicle or corticosterone (∼3.7mg/kg/day), which flattened the endogenous glucocorticoid rhythm. After 7 days, renal arteries were isolated at 7am and 7pm to measure vasoreactivity using wire myography. Other arteries were taken 2-hourly over a 48-hour period for RNA sequencing. In control arteries, endothelium-dependent and independent vasodilation was elevated at wake-phase compared to the sleep-phase. This temporal variation was absent in the renal arteries from corticosterone treated mice. Using CircaCompare and LimoRhyde, we found circadian rhythms in 459 of the 14 225 protein-coding transcripts in control arteries. Following corticosterone treatment, 156 genes lost their rhythm, including canonical clock genes, genes involved in “peroxide homeostasis” such as *Nox4* and “TNF signalling” like *Mmp14*. Paradoxically 492 genes gained a rhythm and were predominantly mitochondrial-related.

**Conclusions:** This study expands the molecular landscape for understanding circadian vascular physiology and emphasises the impact of glucocorticoid rhythm on temporal changes in gene expression and vascular function. This is clinically relevant to the pathogenesis of vascular dysfunction associated with perturbed glucocorticoid signalling for e.g. in metabolic syndrome and chronic stress.

**TRANSLATIONAL PERSPECTIVE:** Disruption of the normal circadian rhythm of plasma glucocorticoid levels causes non-dipping blood pressure, a profile associated with enhanced cardiovascular risk. The role of vascular dysfunction in this context is unknown. We report that arrhythmic plasma glucocorticoid levels impair time-dependent vasoreactivity and alters the circadian transcriptome of the renal artery. We provide a molecular framework for investigations into glucocorticoid-induced vascular dysfunction across the 24-hour day. Clinically, this is relevant for patients on chronic glucocorticoid therapy but could have generalisable implications for the underlying mechanisms of blood pressure rhythm. Targeting rhythm-sensitive vascular pathways may offer novel avenues to improve blood pressure rhythms.

**GRAPHICAL ABSTRACT:** 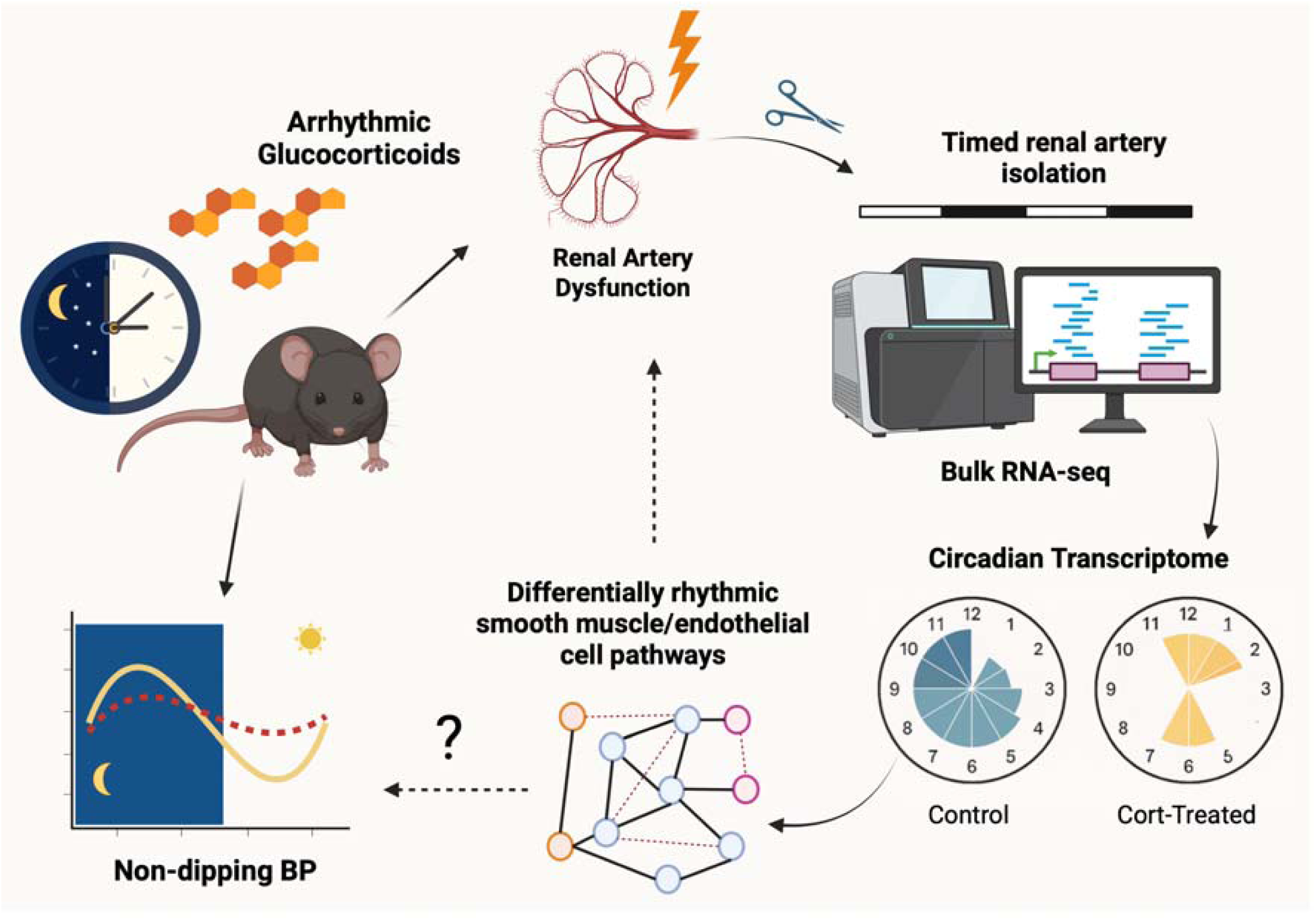

## 1. INTRODUCTION

In health the cardiovascular system exhibits circadian rhythmicity with interrelated parameters, including heart rate, peripheral vascular resistance and blood pressure (BP), oscillating robustly across the ∼24-hour day. BP has a nocturnal “dip”, dropping 10-20% during the sleep period. Attenuation or loss of this profile is associated with adverse cardiovascular outcomes like myocardial infarction and stroke (reviewed in (1)) and can precede and predict disease progression in diabetes (2), kidney disease and in kidney transplant recipients (3). Circadian disruption can also increase the risk for metabolic, psychiatric and immune disorders (4). Resolving circadian disruption may offer a route to improve health but a paucity of information on the molecular framework underlying cardiovascular rhythmicity remains a significant roadblock.

Virtually all cells have an autonomous molecular clock which sets the transcription and translation patterning of the cell’s rhythmic function. These clocks are transcription-translation feedback loops in which positive arm genes, *Bmal1* and *Clock* dimerise to promote the transcription of the negative arm genes *Period* and *Cryptochrome*. The negative arm genes dimerise and translocate to the nucleus, autoinhibiting transcription to reset the cycle in a process taking ∼24 hours. Additionally, the core clock genes control the rhythmic expression of “clock-controlled genes”, and it is estimated that half of all protein-coding genes exhibit circadian rhythmicity in at least one tissue (5). Circadian orchestration is achieved by the master clock in the suprachiasmatic nucleus in the brain, which uses entrainment cues to synchronize internal peripheral clocks with each other and with the external environment. In this context, glucocorticoids are major entrainment factors: circulating levels are strongly rhythmic and, via activation of glucocorticoid receptor (GR), glucocorticoids can adjust the timing of molecular clocks in the liver, lung, cornea and kidney (6). Notably, synthetic glucocorticoids are widely prescribed and the negative impact on circadian physiology may contribute to the increased cardiovascular risk of sustained steroid therapy (7).

The influence of clock genes on cardiovascular function has been explored using knockout approaches. Deletion of any of the core genes induces circadian BP abnormalities, including non-dipping (reviewed in (8)). The cause is unclear but renal salt excretion (9) and peripheral vascular resistance (10, 11) both show circadian rhythmicity. For example, isolated renal artery, mesenteric artery and aorta all show time-of-day variation in response to vasodilating and vasoconstricting agents. Overall, this is characterised by increased reactivity to vasoconstrictors like phenylephrine (PE) during the rest phase (12) and increased reactivity to vasodilators like the endothelium-dependent acetylcholine (ACh) (13, 14) and endothelial-independent vasodilators like sodium nitroprusside (14) during the active period. Similarly, myogenic tone of cremaster artery peaks during the rest phase and troughs during the active phase; a rhythm that is absent smooth muscle-specific *Bmal1* or global *Clock* knock out mice (15).

Both smooth muscle specific (16) and endothelial cell specific (17) deletion of *Bmal1* causes a reduction of BP and a dampening of its circadian rhythm. However, beyond these focussed gene-targeting studies only two microarray studies, both of the mouse aorta, have examined transcriptional oscillations more broadly, reporting rhythmicity of core clock genes and of ∼4% of the protein-coding transcriptome (18, 19). As a conduit artery, the aorta does not contribute substantially to total peripheral resistance, and it is likely that different arterial beds exhibit heterogeneity in their clock-controlled gene expression. The current study focusses on the renal arteries, which strongly autoregulate to control perfusion, glomerular filtration rate and filtered sodium load. Renal haemodynamics contribute importantly to long-term BP stability, and vascular abnormalities may be an early hallmark of kidney disease (20). We have recently shown that chronic glucocorticoid excess induces vasodysfunction in the renal artery, associated with disrupted diurnal rhythm of sodium excretion and induction of non-dipping BP (21). In the current study, we examine the impact of impaired glucocorticoid rhythmicity on time-of-day dependent vasomotor responses in isolated renal artery. We also use RNA sequencing technology to define the impact of impaired glucocorticoid rhythmicity on the circadian transcriptome.

## MATERIALS AND METHODS

### 2.1. Animals

C57BL6J/Crl male and female mice (Charles River, UK), aged 8 to 10 weeks, were used in the study. The mice were housed under a 12-hour light/dark cycle for at least two weeks prior to the experiments, with unrestricted access to water and standard chow (maintenance diet 1; Special Diet Services, Essex, United Kingdom). The lights were turned on at 7:00 AM local time, designated as Zeitgeber time 0 (ZT0), and turned off at 7:00 PM (ZT12). The experiments were conducted under a license from the UK Home Office and with the approval of the University’s veterinary services.

### 2.2. Light Cycle Reversal

To achieve simultaneous access to mice experiencing the subjective light and dark phase, mice were split in 2 cohorts with the second cohort entrained to a reversed light cycle. These mice were acclimatised under 12 hour shifted light conditions for a minimum of 14 days. The control group retained a normal cycle, with lights on at 7am; the reversed group had lights on from 7pm-7am.

### 2.3. Chronic Corticosterone Infusion

Corticosterone was encapsulated in elastomer pellets for gradual release. The in vitro release rate was approximately 3.7 mg/kg per day, as previously described (22), sustained over a period of 4 weeks. The pellets were implanted subcutaneously under isoflurane anaesthesia with buprenorphine analgesia administered (Vetergesic, 0.1 mg/kg subcutaneously). Animals were culled by cervical dislocation for tissue collection 7 days later.

### 2.4. Quantitative PCR

A liver sample was snap-frozen from each experimental animal for quality control assessments. RNA was isolated using the RNeasy-mini kit (Qiagen, US), quantified by the NanoDrop-1000 (Thermo Fisher Scientific, UK) and 500ng cDNA was generated using high-capacity RNA-to-cDNA kit (Thermo Fisher Scientific, UK). Liver abundance of *Arntl*, *Nr1d1*, *Per1* and *Fkbp5* was measured by quantitative RT-PCR using the Universal Probe Library (Roche, UK) Triplicates of each sample and standard curve were run on the LightCycler 480 (Roche, UK). Expression was normalized to the mean concentration of the housekeeping gene Actb on which corticosterone treatment and timepoint had no effect.

### 2.5. Plasma corticosterone measurement

Plasma corticosterone concentration was measured in duplicate using a corticosterone ELISA kit (ADI-900-097; Enzo Life Sciences, UK), which has a detection limit of 27 pg/ml. Following the manufacturer’s protocol, plasma corticosterone was extracted by mixing 10 μL of plasma with 10 μL of diluted steroid displacement reagent (SDR; 1:100 dilution with deionized water) in sterile glass tubes and incubating at room temperature for 5 minutes. Then, 480 μL of assay buffer 15 (AB15) was added to achieve a final plasma dilution of 1:50. In a fume hood, 3 mL of ethyl acetate (Sigma) was added to the glass tubes, which were then thoroughly vortexed. The clear upper organic layer was transferred to a new glass tube and the ethyl acetate was evaporated under nitrogen at 60°C. Samples and standards were prepared and assayed according to the manufacturer’s instructions. Optical density measurements were taken at 405 nm using a microplate reader (Magellan, TECAN), with corrections made at 550 nm.

### 2.6. Ex-vivo circadian vascular reactivity

Two groups of male or female C57BL6/J mice were utilized following either a standard or reversed light cycle as previously described. Each group received either a subdermal corticosterone elastomer pellet or a control pellet. All treatment groups for each time point were assessed concurrently. After euthanasia by cervical dislocation, arteries were quickly dissected, immersed in physiological salt solution (PSS; containing 119.0 mM NaCl, 4.7 mM KCl, 2.5 mM CaCl₂, 1.17 mM MgSO₄, 25.0 mM NaHCO₃, 1.81 mM K₂EDTA, and 5.5 mM D-glucose), and cleaned of attached perivascular fat. A wire myograph system (DMT, Denmark) was used to measure the reactivity of second-order mesenteric arteries and renal arteries. The arteries were allowed to equilibrate for 30-40 minutes under passive tension (at 37°C in a mixture of 95% O₂ and 5% CO₂), and the DMT/Lab Chart Normalisation Module was used to determine and set the optimal pre-tension by calculating the internal circumference (in μm). Maximal contraction was assessed at the beginning and end of the experiment through KCl-induced depolarization (KCl 125mM). Cumulative concentration-response curves for phenylephrine, acetylcholine, and sodium nitroprusside were recorded, with 30-minute washout intervals between each. Responses to phenylephrine were expressed as a percentage of KCl-induced contraction, while vasodilatory responses were expressed as a percentage of phenylephrine-induced pre-constriction.

### 2.7. Circadian bulk-RNA sequencing for renal arteries

48 mice were implanted with corticosterone subcutaneous pellets to manipulate the corticosterone rhythm and 48 were given a vehicle pellet and used as controls. Serial renal artery collection was undertaken every 2 hours for 48 hours which allowed coverage of 2 daily cycles. 4 renal arteries were pooled from 2 mice for RNA extraction. Careful consideration was given to both the frequency of sampling and the number of samples taken at each time point. To obtain sufficient high-quality RNA for RNA sequencing at each time point, the left and right renal arteries from 2 mice were pooled to give n=1. A sample size of n=2 (i.e. 4 mice) was used with collection every 2 hours, as recommended for robust circadian rhythm detection (23, 24). RNA integrity was measured using Agilent Bioanalyzer and an RNA integrity score of 7 was considered sufficient quality to take forward. Sequencing was performed using the Illumina Novaseq Stranded mRNA ligation method for short reads (Edinburgh Genomics). Sequencing data were checked for quality using fast.qc files generated during the sequencing. Adaptors were trimmed using cutadapt version 1.18 with parameters “-a AGATCGGAAGAGC -q 30 -m 46”. The reads were then mapped to the mouse genome (build GRCm39 from Ensembl version 104) using STAR (version 2.7.3a). Expression of each gene was quantified using featurecounts (version 1.5.1) with parameters “-p -t exon -g gene_id -s 2 -Q 10”. Principle component analysis was used in the first instance to assess the variation within the 48 samples. In one sample, the top 20 most highly expressed genes included transcripts (e.g. *Alb*, *Apoa1*, *Apoa2* and *Mup3*) that are uncommon in vascular tissue and were not detected in the other sample taken at the same time point. We interpreted this as being strongly suggestive of contamination of the sample with adipose tissue. This sample (Veh1_15) was excluded from the rhythmicity detection pipeline.

### 2.8. Data Analysis

#### Rhythmicity Detection Pipeline

The bioinformatics tools LimoRhyde (25) and CircaCompare (26) were used to analyse circadian rhythms in RNA-seq data, each serving complementary functions in rhythm detection and comparison. LimoRhyde is a flexible approach based on cosinor regression, which models circadian rhythms by decomposing time into sine and cosine components. It can detect rhythmicity and assesses for differential rhythmicity across conditions, making it particularly useful for datasets that include multiple conditions or more complex experimental designs. By using linear models and empirical Bayes methods, LimoRhyde detects changes in rhythm, amplitude and phase, along with differential expression, offering a robust and statistically rigorous framework for circadian transcriptome analysis. One of its strengths is its ability to handle continuous variables, such as treatment, age or time, and its integration with limma for differential expression analysis. While LimoRhyde is more stringent in detecting differential rhythmicity, reducing the risk of false positives, it can be complemented with other methods for estimating specific rhythmic characteristics, such as amplitude or phase.

CircaCompare, is designed to compare the specific rhythmic parameters—MESOR, amplitude, and phase—between two circadian rhythms. This tool uses a modified cosinor model to perform a direct statistical comparison of rhythmic characteristics between datasets. Unlike LimoRhyde, which primarily focuses on detecting differential rhythmicity and expression, CircaCompare quantifies and tests differences in rhythm characteristics, making it ideal for comparing two distinct circadian rhythms. It provides precise estimates for amplitude, phase, and MESOR differences, which are essential for understanding the biological implications of changes in rhythmicity, such as those induced by different treatments or genetic perturbations. Using both tools in our rhythmicity detection pipeline is beneficial as LimoRhyde offers a comprehensive method for detecting differential rhythmicity across experimental conditions, while CircaCompare enables a detailed comparison of rhythmic parameters between two rhythms. LimoRhyde’s stringent detection ensures robust identification of rhythmic genes, and CircaCompare’s ability to statistically test specific rhythm characteristics provides a deeper understanding of how circadian patterns differ between conditions.

#### Rhythmicity Detection

A raw P-value was calculated to assess the goodness of fit to a cosinor model in a single-condition analysis (control/treatment) with higher P-values indicating a lower likelihood of the gene being circadian. To limit focus to only the most rhythmic genes, we set the P-value cut-off at 0.005 which is more conservative than those used in previous studies (27, 28) which employed P-values of <0.01 and <0.05, respectively. For classification of genes into rhythmic versus non-rhythmic groups P values <0.005 and >0.1 were used.

#### ClusterProfiler

Kyoto Encyclopedia of Genes and Genomes (KEGG) Pathway and Gene Ontology analysis and visualization were done using ClusterProfiler. ClusterProfiler is an R package for gene classification and pathway enrichment analysis, enabling identification of enriched biological themes across gene clusters. It automates the process of analyzing gene sets across multiple species, and allows comparisons across various biological ontologies, such as GO, KEGG, and Disease Ontology. The package integrates classification, enrichment analysis, and visualization into a seamless workflow. It employs a hypergeometric test for enrichment analysis, and controls for multiple testing by using q-values (FDR correction).

## 3. RESULTS

### 3.1. Arrhythmic circulating glucocorticoids attenuated time of day variation in endothelial-dependent and independent vasodilation

In male C57BL6 mice implanted with a slow-release pellet containing corticosterone, plasma corticosterone was elevated compared to the control (vehicle) group and the normal day-night variation was lost (**Figure 1A**). Glucocorticoids have previously been shown to synchronise the rhythm of core-clock genes in the liver (29). We examined this and found that arrhythmic circulating glucocorticoid suppressed the mRNA expression rhythm of the early glucocorticoid responsive gene, *FK506 binding protein 5 (Fkbp5)*, compared to control mice (**Figure 1B**). The rhythms of core clock genes, *Per1* and *Arntl* were also dampened (**Figure 1C-D**) in the liver of corticosterone treated mice.

**Figure 1:**
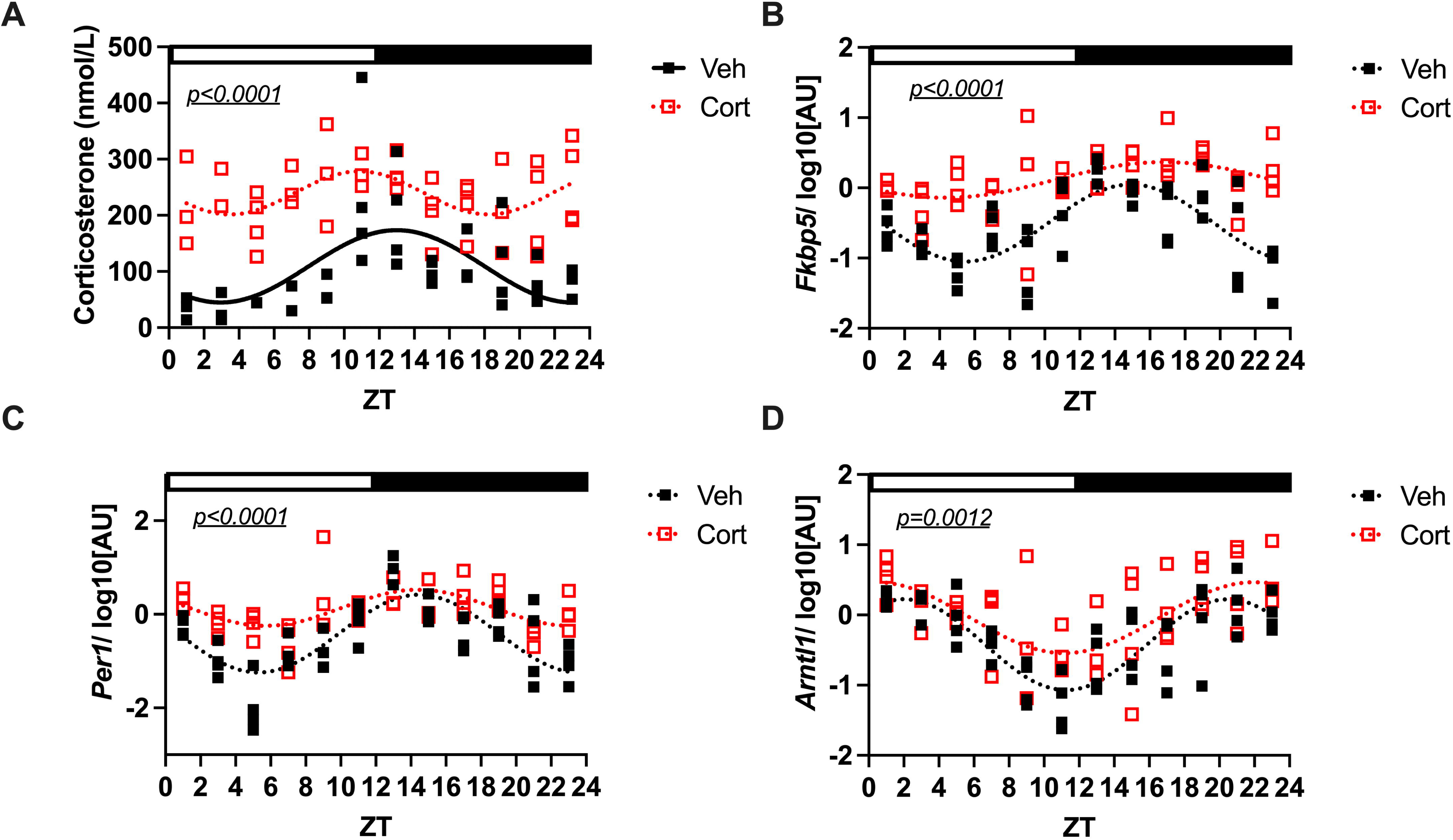
(A) Plasma Corticosterone levels across the 24-hour day, (B) liver *Fkbp5* (C) *Per1* and (D) *Arntl* mRNA expression in male C57BL6J mice treated chronically with vehicle or corticosterone. Mice were treated with a subcutaneously inserted corticosterone or vehicle pellet. After a week, mice were euthanised and plasma and livers sampled at 2-hourly intervals over 48 hours. Individual values are shown with group mean ± SD; n=2-4, statistical comparison is an Extra Sum of Squares test testing the hypothesis that one curve fits both data sets. White and black bars indicate light and dark phases respectively. ZT time is indicated on X axis, where ZT 0 is the time of lights on.

Next, we examined time-of-day dependent vasoreactivity in isolated renal artery. In the control group, the vasodilation response to acetylcholine (ACh), which is endothelium-dependent, exhibited time-of-day dependent variation: sensitivity to ACh was higher in the wake phase (ZT12) compared to the sleep phase (ZT0, **Figure 2A**). The vasodilatory response to the nitric oxide donor SNP, which is independent of the endothelium, also exhibited overt time-of-day variation (**Figure 2E**) and like the response to ACh, sensitivity to SNP-induced dilation was higher during the wake phase (ZT12) compared to sleep phase (**ZT0, Figure 2E**).

**Figure 2:**
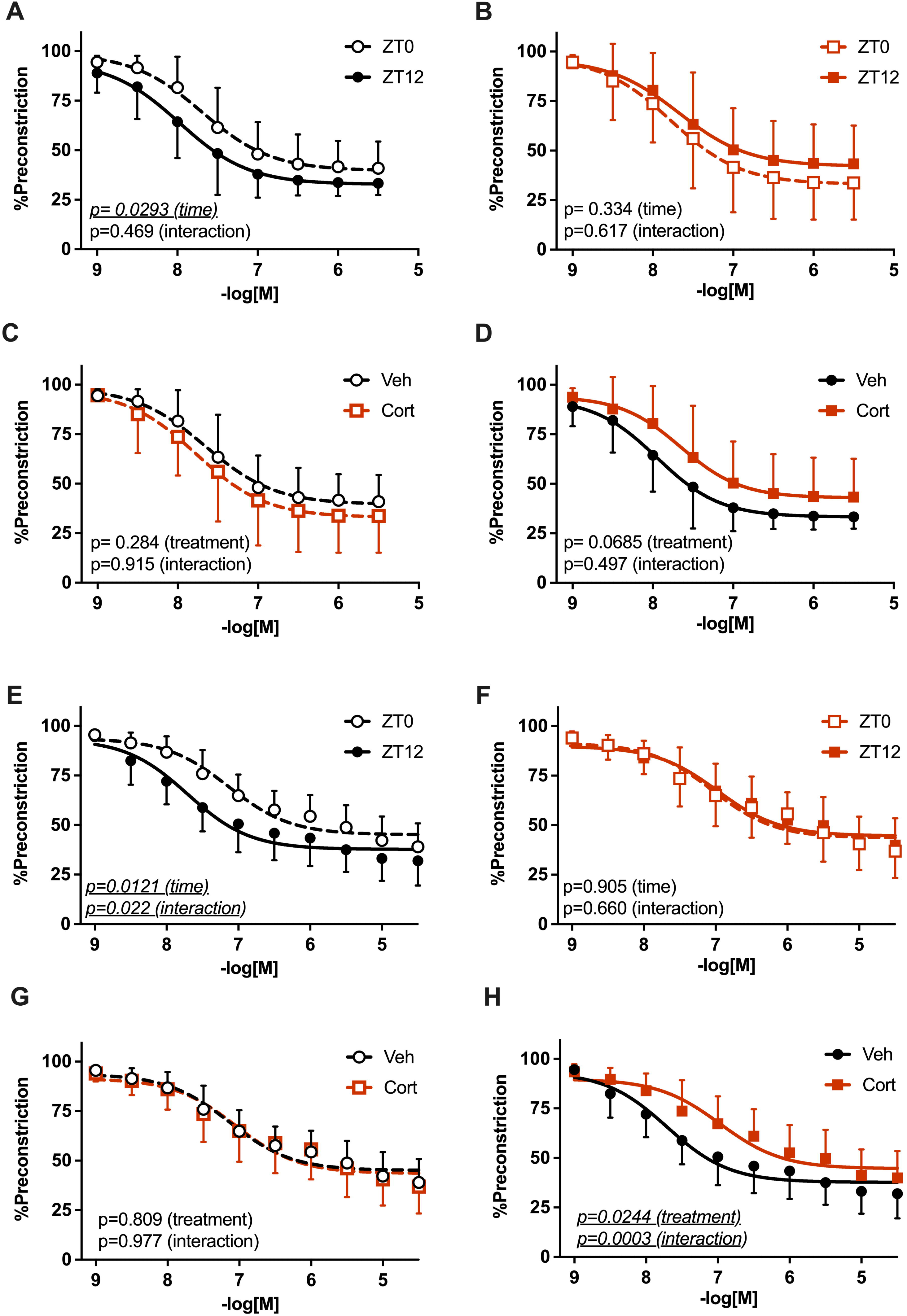
Vasodilatory response to acetylcholine (ACh) and sodium nitroprusside (SNP) exhibits time-of-day dependent variation in vehicle treated control renal arteries but not in renal arteries from corticosterone-treated male mice. Renal arteries were dissected from male mice treated with vehicle (blank silastic pellets) and corticosterone (corticosterone silastic pellets, release rate ∼3.7mg/kg/day) and mounted onto an isometric wire myograph. Arteries were pre-constricted with phenylephrine before being subjected to increasing concentrations of ACh or SNP. The vasodilatory response to ACh was compared during the active phase (ZT12) and inactive phase (ZT0) in renal arteries from vehicle treated mice (A) corticosterone-treated mice (B) and comparison between vasodilatory response to ACh in vehicle (veh) and corticosterone-treated (cort) mice was made during the inactive phase (C) and active phase (D). The vasodilatory response to SNP was compared during the active phase (ZT12) and inactive phase (ZT0) in renal arteries from vehicle treated mice (E) corticosterone-treated mice (F) and comparison between vasodilatory response to SNP in vehicle (veh) and corticosterone-treated (cort) mice was made during the inactive phase (G) and active phase (H). Data are expressed as a % of the pre-constricted value and are mean ± SD and analysed by matched two-way ANOVA, significant p values are underlined and italicized, n=12.

In corticosterone-treated mice, temporal variation in endothelial-dependent vasodilation of the renal artery was absent (**Figure 2B**). Endothelial-dependent vasodilation was comparable to the control arteries during the sleep phase (**Figure 2C**) but exhibited reduced sensitivity in the wake phase (**Figure 2D**). Cort-treated renal arteries exhibited no diurnal variation in response to SNP (**Figure 2F**) with their vasodilation comparable to control arteries during the sleep phase (**ZT0, Figure 2G**) but with markedly reduced sensitivity compared to controls during the wake phase (**ZT12, Figure 2H**).

The contractile response to a depolarising stimulus (125mM KCl) was similar in both experimental groups and did not show time-of-day variation (**Supp Figure 1A**). The vasoconstriction response to PE was unchanged between time points in the control group (**Supp Figure 1B**) but surprisingly, the arteries from corticosterone-treated mice did exhibit time-of-day variation with reduced sensitivity to PE during the sleep phase (**ZT0, Supp Figure 1C**). Renal arteries from corticosterone-treated mice displayed a reduced sensitivity to PE at both time points (**Supp Figure 1D, E**).

### 3.2. The renal artery circadian transcriptome is altered by chronic glucocorticoid treatment

To gain a molecular-level insight into the changes that may underlie circadian renal vascular function, an unbiased approach was taken using bulk RNA sequencing of renal arteries in vehicle and corticosterone-treated mice collected every 2 hours over a 48-hour period. The entire dataset including CircaCompare and LimoRhyde analyses are available to download online (**Supp Table 1**).

In renal arteries from control mice, 16,256 transcripts were detected with 14,225 protein coding genes. Taking only the protein coding transcripts, regression to a cosinor waveform was performed using LimoRhyde and a p-value threshold of p<0.005. This gave 459 rhythmically expressed genes (REGs) which was ∼3% of the total protein coding genes (**Supp Table 2**). In the renal arteries from corticosterone-treated mice 16 725 transcripts were detected, 14 806 of these were protein coding. Using the same criteria as control arteries, 917 of these were REGs, which was ∼6% of the total protein coding transcriptome (**Supp Table 3**). Only 140 of these REGs were common to both vehicle and cort-treatment groups (**Figure 3A, D**) and the known clock genes were contained in this group. These 140 common REGs were further analysed using CircaCompare to assess changes in rhythmic parameters like amplitude or acrophase (time of peak) between the two groups: 20 genes exhibited reduced amplitude in the corticosterone group compared to vehicle, 32 had altered acrophase and 7 genes had reduced amplitude and altered acrophase (these were exclusively clock genes, **Supp Figure 2**, **Supp Table 4**).

**Figure 3:**
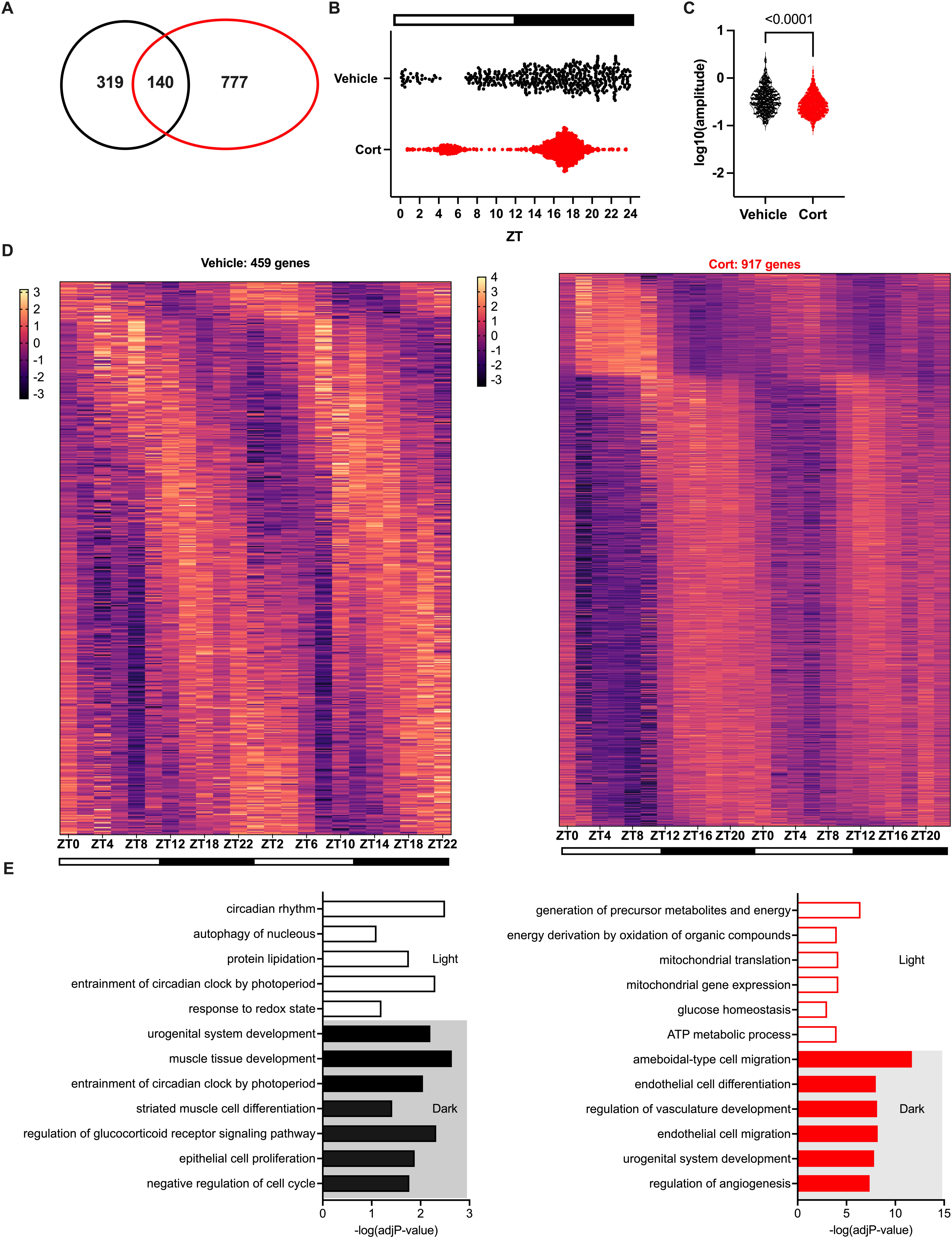
Transcriptional rhythmometry by bulk RNA-sequencing reveals 459 rhythmic transcripts in vehicle versus 919 rhythmic transcripts in renal arteries from corticosterone-treated male C57BL6/J mice. Renal arteries were dissected from male mice treated with vehicle (blank silastic pellets) and corticosterone (25mg, corticosterone silastic pellets) in 2-hourly intervals across the 24-hour day and RNA-sequenced. Transcripts were analysed by LimoRhyde with p value for cosinor regression <0.005. (A) Venn diagram indicating the rhythmic transcripts in vehicle (black-459), corticosterone-treated (red-917) and the 140 transcripts in common between the 2 groups. (B) Dotplot of acrophases as calculated by *CircaCompare* of each of the 459 and 917 rhythmic genes in renal arteries from vehicle or corticosterone-treated mouse renal arteries respectively across the 24-hour day. Each point represents a single transcript’s time of peak expression. (C) Violin plots indicating the logarithmic value of the amplitude of each of the rhythmic transcripts detected in the vehicle and corticosterone-treated renal arteries. (D) Heatmap showing the 459 rhythmic genes defined using LimoRhyde. Genes are plotted as Z scores and by increasing acrophase. Each column represents the z score of each gene at each time point for each biological replicate. Black bars below represent the active period and white, the inactive. (E) Bar plot indicating the most enriched terms the rhythmic transcripts belong to and corresponding to each distinct light phase, according to Gene Ontology Biological Processes and predicted by ClusterProfiler. Light phases are indicated on the diagram and on the X axis the negative logarithm of the adjusted P-value is shown.

Comparing the REGs, markedly different patterns of expression were seen between groups (**Figure 3D**). The acrophases, or times at the peak expression, in the control group were distributed evenly across the 24-hour day with a reduced number of acrophases during the sleep period (**Figure 3B**). Whereas in the corticosterone-treated group, acrophases were confined to two clusters with the majority clustering around ∼5 hours after the lights off time (during the wake phase) and a smaller cluster aligning to ∼5 hours after the lights on time (during the sleep phase, **Figure 3B**). The amplitudes of REGs in the corticosterone-treated group were also lower compared to the amplitudes of REGs in the vehicle group (**Figure 4C**). Gene ontology analysis of REGs in the control group gave enrichment for terms including “circadian rhythm”, “entrainment of circadian clock by photoperiod” and “muscle-tissue development”. In contrast, GO terms related to circadian rhythm were absent from the REGs in the cort-treated group, which was instead enriched for terms including “endothelial cell differentiation”, “endothelial cell migration” and “generation of pre-cursor metabolites and energy” (**Figure 3E**). There were 492 genes that gained de novo rhythms following chronic corticosterone treatment, i.e. they were exclusively rhythmic in the corticosterone-treated group (P<0.005 for corticosterone-treated and P>0.1 for vehicle, **Supp Table 5**). Pearson correlation analysis between the top 20 genes suggest co-expression of many of these genes (**Supp Figure 3**, for e.g. *Krit1* and *Aggf1* R=0.93 and *Nuak1* and *She*, R=0.95). 4/20 of these genes are mitochondrial membrane genes (*Slc22a3*, *Pdss2*, *Ptpmt1*, *Epha4*) and Nuak1 regulates mitochondrial function (30). *She* regulates vascular lumen size (31). Krit1 and Aggf1 polymorphisms both cause vascular malformations (32).

**Figure 4:**
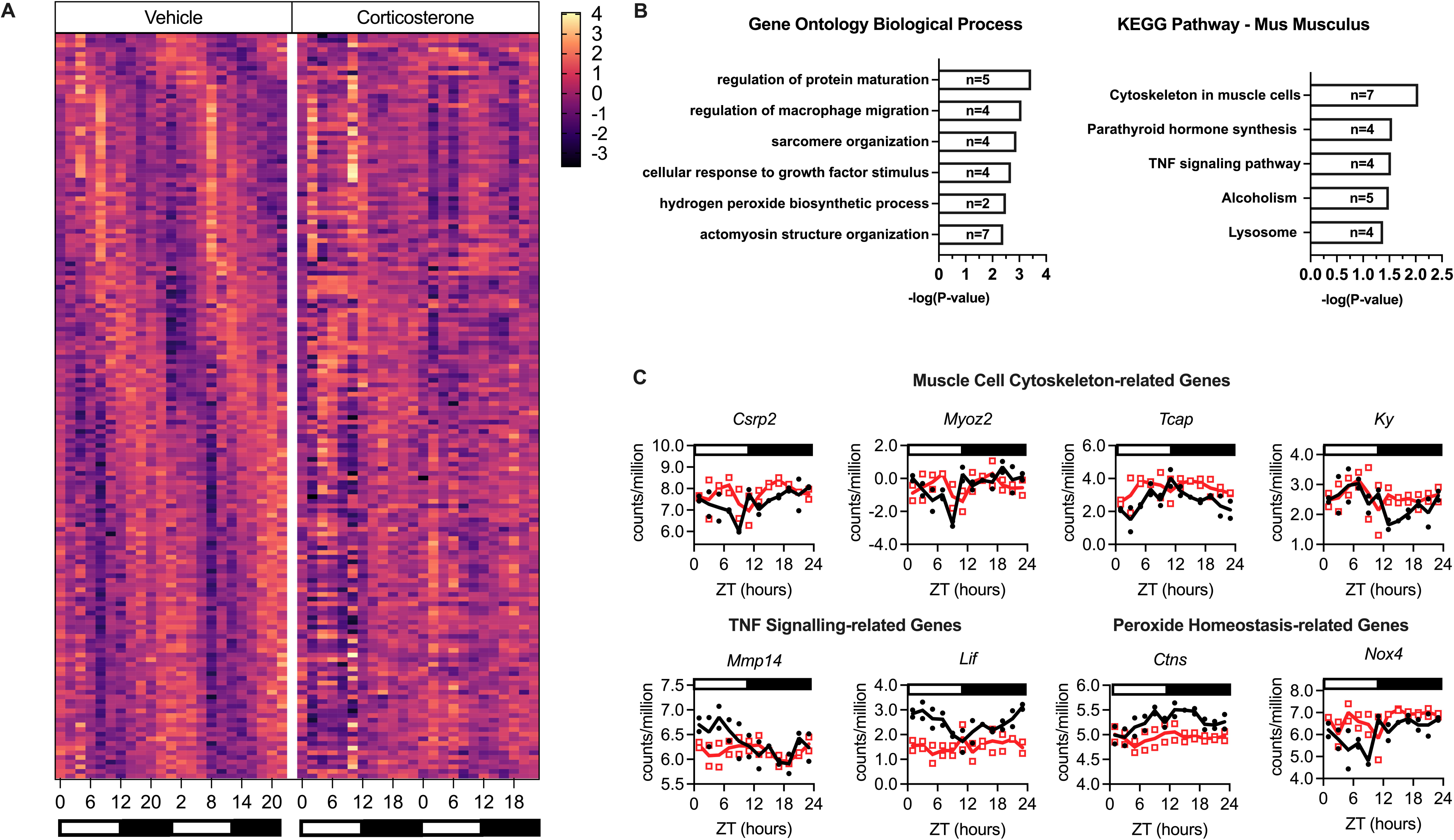
160 transcripts lose rhythmicity in corticosterone-treated renal arteries. (A) Heatmap showing the 160 rhythmically expressed genes (REGs) in the vehicle group that become arrhythmic in the corticosterone-treated group. Genes are plotted as Z scores and by increasing acrophase. Each column represents the Z score of each gene at each time point for each biological replicate. Black bars below represent the active period and white bars the inactive. (B) Most significant GO-terms the arrhythmic transcripts belong to according to Biological Processes and Kyoto Encyclopaedia of Genes and Genomes as predicted by ClusterProfiler Gene expression profiles in vehicle and corticosterone-treated renal arteries (shown as logarithmic counts per million). (C) Gene profile examples of REGS exclusive to vehicle treated mice most enriched within the muscle cell cytoskeleton, peroxide homeostasis and TNF signalling pathways. White and black bars indicate the inactive (light) and active (dark) phases respectively and each dot represents an individual biological sample (n=2 where each n is a mix of both renal arteries from 2 mice. For scatter graphs, the vehicle treated control group is represented by black annotation with corticosterone treated group red. ZT time is indicated on X axis, where ZT 0 is the time of lights on.

### 3.3. Arrhythmic plasma glucocorticoids caused the loss of rhythm of 156 genes, downregulated 202 genes and upregulated only 9 genes

We anticipated that the genes with rhythms abolished by corticosterone treatment would be most relevant to the cort-induced changes in temporal renal artery function and non-dipping blood pressure (22). Using our stringent thresholding for rhythmicity (P<0.005 in vehicle group, P>0.1 in the cort group), 33% of the vehicle REGs were exclusively rhythmic in the vehicle group (156 out of the 459 rhythmic genes) having lost their rhythm following cort-treatment (**Figure 4**, **Supp Table 6**). These included genes involved in “muscle cell cytoskeleton” like *Ky* and *Csrp2*, “peroxide homeostasis” like *Ctns* and *Nox4* and “TNF-signalling” like *Mmp12* and *Lif* (Figure 4B-C).

We next performed a differential expression analysis to investigate changes in renal artery genes in response to corticosterone treatment irrespective of time. This approach favours genes with limited variation throughout the day. Using a log fold change of 1.2 and False discovery rate (FDR) of 0.01, we found 211 differentially expressed genes (**Figure 5**, **Supp Table 7**). 202 of these were downregulated in response to cort treatment with 9 genes upregulated. Downregulated genes included genes involved in local tissue corticosterone steroidogenesis including *Cyp11b1* which catalyses the conversion of 11-deoxycorticosterone to corticosterone and *Hsd11b2* which deactivates corticosterone (converting it from corticosterone to deoxycorticosterone), genes involved in apelin signalling: both the apelin receptor (*Aplnr*) and apelin protein (*Apln*), and extracellular matrix genes, including *Col6a5* and *Col28a1*. The upregulated genes included *Timp4* and *Ltf*. *Timp4* encodes a peptidase involved in degradation of extracellular matrix and control of smooth muscle cell proliferation (33). *Ltf* encodes the lactoferrin, which has immunomodulatory functions in arteries and is implicated in the progression of atherosclerosis (34).

**Figure 5:**
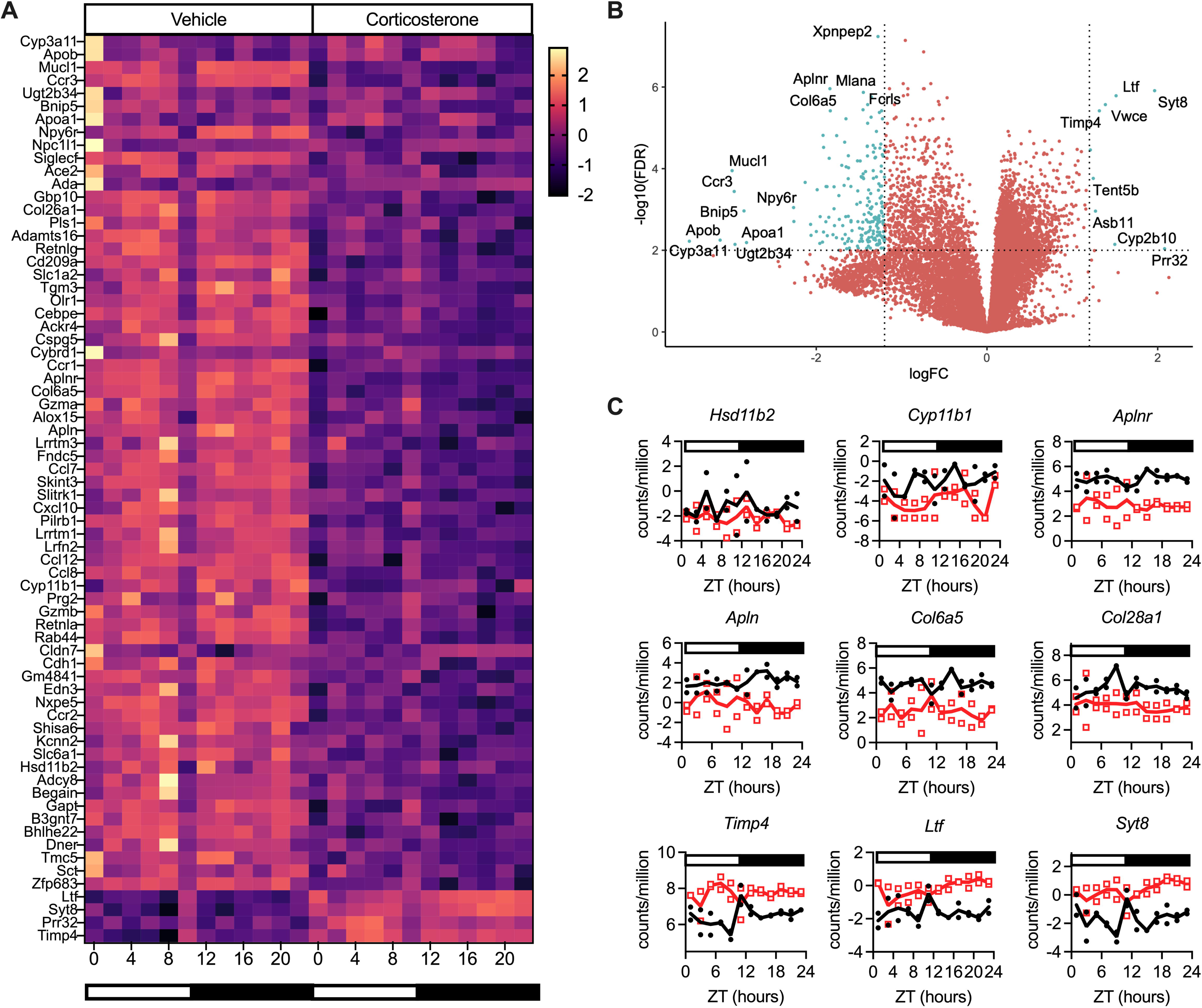
Differential expression analysis of transcripts most changed between renal arteries from vehicle and corticosterone-treated C57BL6/J male mice. (A) Heatmap displaying the top differentially expressed transcripts in renal arteries from vehicle and corticosterone-treated mice. (B) Volcano plot of differentially expressed transcripts under arrhythmic corticosterone this is logFC1.2. (C) Example gene expression profiles of top differentially expressed genes. For scatter graphs, the vehicle treated control group is represented by black annotation with corticosterone treated group red. White and black bars indicate the inactive (light) and active (dark) phases respectively and each dot represents an individual biological sample (n=2 where each n is a mix of both renal arteries from 2 mice). ZT time is indicated on X axis, were ZT 0 is the time of lights on.

### 3.4. Differential rhythmic expression of Vascular Smooth Muscle Specific genes

Our unbiased RNA sequencing approach did not detect any obvious candidate genes underlying the intrinsic functional changes we detected in myography experiments. Next, we focussed on genes involved in vascular smooth muscle physiology which is where we found the biggest differences in vascular function (response to SNP). We used the Hamonizome 3.1 Gene-Phenotype associations gene set for abnormal vascular smooth muscle physiology (35) and tested whether these genes exhibited any changes between corticosterone and vehicle groups. There are 110 genes in this set (16 were not found in our dataset) and we used the CircaCompare analysis for MESOR as this considers the variation across the day. We separated these by those with increased MESOR and those with decreased MESOR (p<0.001 for MESOR difference between control/corticosterone). The top increased genes included *Trpv4*, *Nos3*, *Pecam1*, *Kcna5* and *Slc4a7* (**Figure 6**, **Supp Table 8**). These were elevated during the wake period and the 24-hour pattern of this group of genes strongly correlates (**Figure 6B**), suggesting that these might be co-expressed. Intriguingly, the transcription factor serum response factor (*Srf*), which controls gene transcription in vascular smooth muscle cells, shares a strikingly similar pattern with this group of genes. The difference in expression between corticosterone and control groups was observed during the active period, aligned with the largest functional differences in terms of vascular reactivity. The downregulated genes included *Mmp2*, *Mmp9*, *Col3a1* and the endothelin B receptor (*Ednrb*). Aside from Mmp2 and *Col3a1*, these were not strongly correlated (**Figure 6B**).

**Figure 6:**
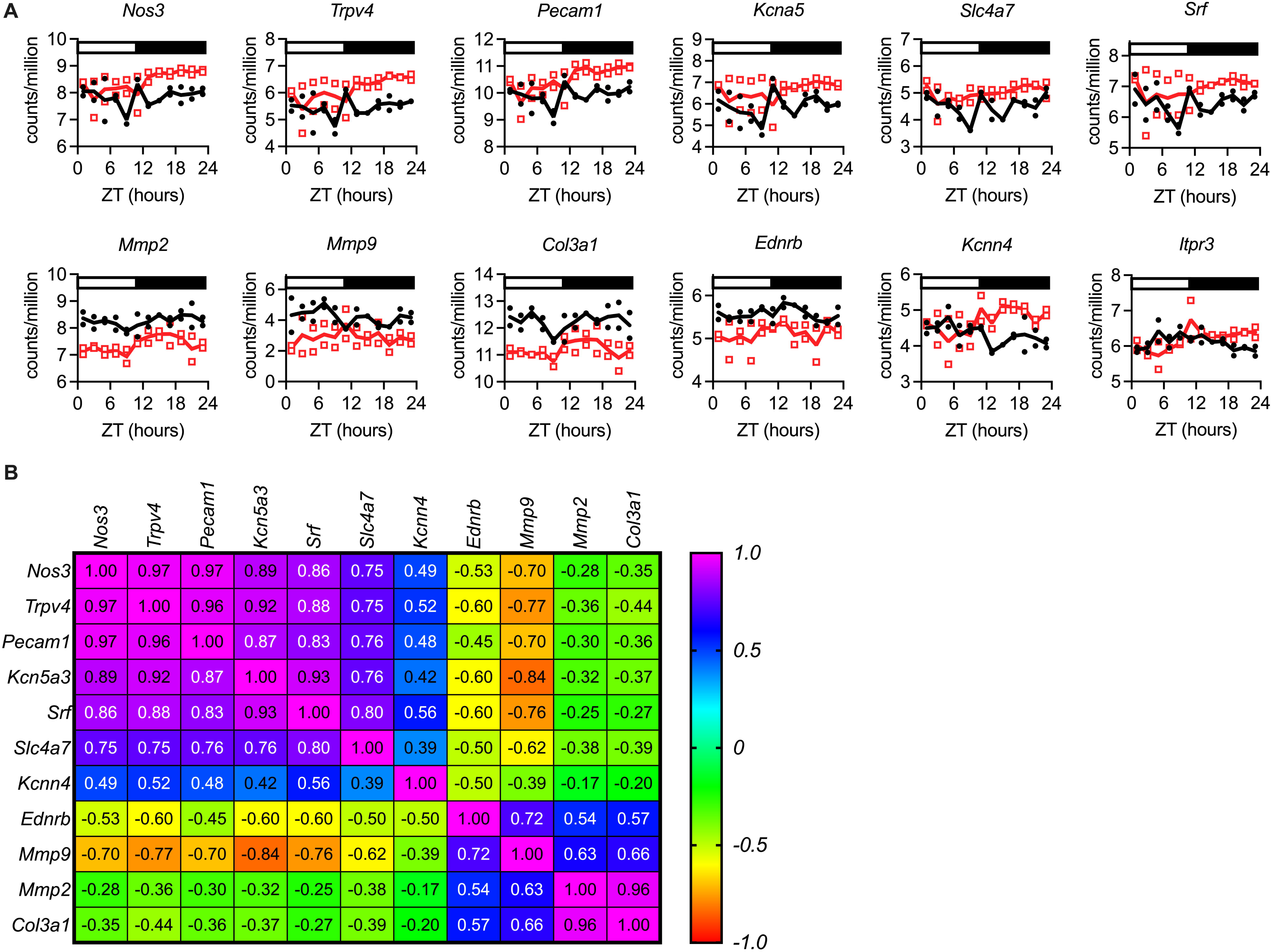
Genes from the Vascular Smooth Muscle Gene set with most significant difference in MESOR between renal arteries from vehicle and corticosterone-treated mice. Transcript expression of genes from the vascular smooth muscle gene set from Harmonizome 3.1 was regressed to a cosinor fit by Circacompare and MESOR compared between renal arteries from vehicle and corticosterone treated mice. (A) Example gene expression profiles and (B) Pearson correlation heatmap showing the R-values for correlation between the genes with most significant MESOR differences, where R=1.0 is perfect positive correlation. For scatter graphs, the vehicle treated control group is represented by black annotation with corticosterone treated group red. White and black bars indicate the inactive (light) and active (dark) phases respectively and each dot represents an individual biological sample (n=2 where each n is a mix of both renal arteries from 2 mice). The vehicle treated control group is represented by black annotation with corticosterone treated group red. ZT time is indicated on X axis, were ZT 0 is the time of lights on.

We also investigated genes that exhibited changes in acrophase (according to CircaCompare, p<0.001, **Supp Table 8**), these included only 6 genes *Kcnn4* (p=9.6×10^-7^, peak time 3 for vehicle and 16 for corticosterone), *Itpr3* (p=0.00012, peak vehicle 9, peak for corticosterone 15) *Chrm3*, the gene for the muscarinic ACh receptor (p= 0.00037, peak 9 for veh and 15 for corticosterone), *Ptger3* (the gene encoding EP3 receptor for prostaglandins, p=0.003, peak vehicle 14, peak corticosterone 5).

## 4. DISCUSSION

Glucocorticoids are widely prescribed but chronic therapy has numerous side-effects, including a substantially increased cardiovascular risk (36). Oral glucocorticoid therapy can disrupt the diurnal rhythm of endogenous cortisol (37). This induces circadian disruption in key cardiovascular systems, such as the kidney (22) but this may extend to the arterial vasculature too. The current study provides a detailed description of the circadian transcriptome of the renal artery and highlights the extensive disruption induced by sustained exposure to arrhythmic circulating glucocorticoid.

We collected arteries at 2-hourly time points and used LimoRhyde (25) and Circacompare (26) in a rhythmicity detection pipeline. These algorithms offer complementary analysis. LimoRhyde provides a comprehensive method for detecting differential rhythmicity across experimental conditions, while CircaCompare enables a detailed comparison of rhythmic parameters between two rhythms. We confirmed that under control conditions the renal artery expresses all canonical circadian clock genes and their paralogues, and these oscillate tightly and robustly. We found that ∼3% of the protein-coding transcriptome was rhythmic, which approximates that in the aorta (4%, (19)). The slight discrepancy may be biological or may reflect different transcriptomic technologies (RNA sequencing versus microarray). We also implemented a more stringent algorithm, lower p value threshold and increased sampling frequency. Nevertheless, the overall conclusion is that the aorta (conduit artery) and renal artery (hybrid conduit and resistance) are less intrinsically rhythmic, having fewer rhythmically expressed genes than tissues such as the kidney at 13% and the liver at 16% (5). This suggests that arteries are more reliant on reactive homeostatic adaptation than predictive circadian adaptation. Consistent with this interpretation, a recent study reported in skeletal resistance arteries that only ∼25% of myogenic tone was rhythmic with 75% of the total myogenic tone being non-rhythmic and constitutively present (15). It is also possible that some tissues might only need a small number of REGs to orchestrate functional rhythmicity at the protein level (e.g. phosphorylation and other post-translational modifications)(38).

### 4.1. Arrhythmic plasma glucocorticoid levels induce *de novo* rhythms

Glucocorticoids have a prominent role as peripheral clock entrainment factors (6) and exposure to sustained but arrhythmic corticosterone caused overt dampening of core circadian clock genes. This was characterised predominantly by a reduction in amplitude of the molecular clock genes, but some transcripts also showed changes in acrophase (figure x). A similar finding has been reported in skeletal muscle (39). Surprisingly, the loss of rhythmicity or mis-aligned peak expression of the core clock, did not lead to a large-scale reduction in the number of rhythmically expressed genes. In fact, the corticosterone treated group exhibited double the number of REGS compared to the controls. Examining these REGS in more detail, we found the acrophases mostly occurred at either ZT 5 and ZT17, ∼5 hours after the environmental light: dark and dark: light transitions. We propose that the expression of such genes is driven by light-induced transcription, although this could be secondary to light-driven behavioural changes (40). There is evidence that glucocorticoids cause an increase in arousal (41) and out of phase glucocorticoid levels causes changes in feeding behaviour in rats (42). This was not assessed in the current study but previously we found that chronic glucocorticoid did not affect locomotor activity, feeding behaviours or heart rate (22). Nevertheless, this interpretation was based on hourly measurements of x-y movement by radiotelemetry and on assessment of 12-hour food intake, so it is possible that more subtle activity/feeding changes may have been overlooked.

A second, more speculative, possibility is that *de novo* expression rhythms induced by glucocorticoid exposure, reflect the perturbed hemodynamic environment of the renal artery. During periods of sleep, when BP falls, arteries can experience altered shear stress compared to the wake period. Chronic corticosterone infusion induces non-dipping BP, and it would be anticipated that shear stress may be more similar to wake period levels. Many vascular genes are regulated by shear stress, including *KCNN3* (43) and *KCNN4* (43, 44) which are rhythmic only in the corticosterone group, peaking at ZT17 and ZT16 respectively. These encode small-conductance calcium-activated potassium channels and act to hyperpolarise the endothelium to induce vasodilation. Adding to this concept, the GO terms enriched within rhythmic genes in the corticosterone-treated group includes “endothelial cell proliferation” and “mitochondrial genes”; increases in shear stress are known to promote endothelial adaptation (44) to new mechanical demands and changes in mitochondrial genes may fuel these new energetic needs.

### 4.2. Glucocorticoids cause attenuation of the normal vasodilation in the active period

We found an enhanced vasodilatory response to ACh during the active period, consistent with other studies in aorta (14) and mesenteric arteries (13). This circadian behaviour of elevated active-phase vasodilation is thought to offset higher sympathetic output in the active phase, thereby buffering BP (15). ACh activates muscarinic ACh receptors on the endothelium, and the encoding gene *Chrm3* was rhythmic in our control group, peaking just prior to the active period. The activation of endothelial Chrm3 increases the activity of the endothelial nitric oxide synthase (eNOS, encoded by *Nos3*) to increase endothelial derived nitric oxide production. In humans the acetylcholine-induced elevation in forearm blood flow peaks during the active period (45). Others show Nos3 mRNA and protein in mesenteric arteries is elevated during the active phase (13) and enzymatic activity is rhythmic in the aorta (46). Notably, we did not find a transcriptional rhythm in *Nos3* in the renal artery from control mice.

We also found elevation in the renal artery vasodilatory response to SNP. SNP is a NO donor and bypasses the Chrm3-eNOS pathway in the endothelium, acting directly on smooth muscle soluble guanylyl cyclase, to increase the formation of cyclic guanosine monophosphate (cGMP). cGMP induces smooth muscle relaxation by protein kinase G-induced reduction of intracellular calcium through IP3 signalling pathways, SERCA and calcium activated K channels. Formation of sGMP exhibits a daily rhythm in rat aorta with a peak late in the inactive phase (47). There is no evidence in our dataset that any of the paralogues of guanylyl cyclase exhibit transcriptional rhythms. Studying protein and post-translational changes would be more informative here.

Our week-long corticosterone treatment abolished the temporal difference in vasodilatory response to both ACh and SNP. This manifests as a reduced vasodilatory response to SNP and a trend for a similarly reduced vasodilatory response to ACh during the active period. *Chrm3* remains rhythmic in the corticosterone group, but peak expression was significantly phase shifted, moving 6 hours to the middle of the active phase. Glucocorticoid treatment also increased *Nos3* expression, evident only in the active period. However, the functional significance is not clear because we find endothelial-dependent vasodilation of the renal artery, which is Nos3-dependent, tends to be reduced at this time point. Indeed, glucocorticoids reduce circulating nitic oxide bioavailability (48). This is largely reflective of iNOS inhibition (48) and in our study the encoding gene, *Nos2*, was downregulated in corticosterone treated mice.

That glucocorticoid-treatment significantly reduced SNP-dependent vasodilation is suggestive of an inhibitory effect on the pathway connecting guanylyl cyclase mediated production of cGMP, protein kinase G-dependent phosphorylation of calcium activated K channels (e.g. Kcnn3 and Kcnn4), and inhibition of IP3 signalling and elevated SERCA. Others have shown that glucocorticoids suppress the activity of guanylyl cyclase in rat vascular smooth muscle cells (49). They also can affect cGMP degradation by reducing the activity of phosphodiesterase 5, PDE5 (50). In our study, corticosterone did not affect guanylyl cyclase transcription. However, *PDE5A*, which was not rhythmically expressed, was elevated by corticosterone-treatment, which would cause specific degradation of cGMP. We also saw group differences for *Itpr3* expression, which encodes the receptor for IP3.

Expression was rhythmic in both groups but with significantly different acrophases (ZT 9 for control and ZT 15 for corticosterone). Binding of IP3 to its receptor results in more intracellular calcium release and contraction. If receptor level follows transcription, we might expect a greater susceptibility for contraction in the late active period in the corticosterone group and early active period for the vehicle group.

Another important component of redox signalling in vasculature is the endothelial derived hyperpolarising factor, H_2_O_2_, constitutively generated by *Nox4*, a NADPH oxidase expressed in all vascular cell types (51). *Nox4* is regulated at the gene transcription level, and we found rhythmic expression in controls with a peak at the start of the active period and trough during rest. This is consistent with its known circadian rhythm, with its promoter activated by the core clock (52) and fits with the idea of enhanced active-phase vasodilation. *Nox4* expression rhythm was strikingly absent in the corticosterone group, reflecting sustained elevation of expression throughout the rest period. Our bulk RNAseq does not allow us to resolve the cell type and *Nox4* is expressed in both endothelial and smooth muscle cells. H_2_O_2_ has vasodilatory actions in endothelium (39) but contractile in the smooth muscle (reviewed in (53)). As this is bulk RNA expression data from renal artery containing both smooth muscle and endothelium we cannot ascertain where the Nox4 expression may be localised. Sustained daytime Nox4 levels in the corticosterone-treated group in the endothelium may be a compensation mechanism, or if the expression is smooth muscle in origin, it may be the cause of the elevated daytime BP. Further scrutiny of the cellular localisation of Nox4 and its role in the circadian rhythm of BP is warranted. *Nox4* also has a role in regulating arterial stiffness; elevated mitochondrial *Nox4* expression increased aortic stiffness and reduced the response to phenylephrine in aorta (54). This is intriguing as we also see reduced phenylephrine-induced contraction in corticosterone treated groups at both time points.

### 4.3. Glucocorticoids cause changes to genes involved in vascular morphology

The other broad theme that emerges in our data is glucocorticoid-induced changes to genes involved in vascular morphology including changes to genes involved in the muscle cell cytoskeleton and extra-cellular matrix (ECM) regulation. ECM is a dynamic and constantly remodelling structure, and regulators are known to have a circadian rhythm heavily influenced by the molecular clock (55). The rhythms of *Mmp14* and *Adamts4* are well described in other tissues with Mmp14 rhythm also described in the glomerulus (56). These generally peak during the rest phase; we report similar rhythms in renal artery under control conditions and we see attenuation with corticosterone treatment, with flattening of the *Mmp14* rhythm characterised by reduced rest phase expression and reduced MESOR of *Adamts4*. Beyond circadian rhythm analyses, in our differential expression we see suggestions of ECM remodelling with markedly reduced collagen genes (*Col3a1*, *Col28a1*, *Col6a5*, *Col26a1*) but elevation of *Timp4*. TIMP proteins inhibit metalloproteinases like MMP14, which cause proteolysis of the ECM and increased turnover, therefore elevated TIMPs usually suggests increased ECM accumulation. In carotid artery injury there is increased TIMP4 levels and increased collagen deposition (57). It is important to note that these gene changes occur following a single week of arrhythmic glucocorticoid levels. We might expect to see further signs of vascular remodelling with longer term exposure. In a mouse model of Cushing Syndrome for e.g. renal artery changes to ECM include downregulation of *Elastin*, *Emilin1* and upregulation of *TGFß* (21). We did not see any of these changes in renal arteries exposed to short-term arrhythmic glucocorticoid.

### 4.4. Limitations

This study has several limitations. First, we carried this experiment out under normal light: dark conditions, this was intentional to keep the experiment consistent with the environment experienced by mice undergoing blood pressure monitoring previously (22) and is more reflective of the human experience of light: dark exposure. However, it makes it impossible to separate gene changes as the result of intrinsic circadian rhythms versus changes driven by behaviour that follows the light-dark cycle. Second, the data is derived from bulk renal arteries and is therefore heterogeneous with both smooth muscle and endothelial derived cells. It is not possible to assign the gene changes seen to specific cells. Thirdly, we only carried out this work using male mice, but others have shown sexual dimorphism in BP control in circadian clock knock out mice (58) and testosterone itself can influence Nox4 expression (59).

### 4.5. Conclusion and Perspectives

This study demonstrates that hundreds of genes in the renal artery oscillate with a 24h period. This, and the diurnal variation in the vasodilatory function of the renal artery, might contribute to time-of day variation in renal blood flow and glomerular filtration rate. This nuanced control of renal haemodynamics serves a vital function underpinning organ metabolism and systemic electrolyte balance. Importantly, we found that glucocorticoids are significant circadian disruptors of renal artery physiology, both at the transcriptional and functional levels. The long-term consequences of the loss of rhythmicity are unknown but may contribute to the adverse effects of glucocorticoid excess on kidney and cardiovascular physiology (60). Our data set provides a valuable resource for the cardiovascular research community, enabling a deeper understanding of circadian rhythmicity of pathways important for vascular structure and function. The temporal nature and sampling frequency of the data also lends itself to correlation analysis, with tools like Graphia (61), which may be another route for in depth analysis of specific vascular pathways of interest.

## ACKNOWLEDGMENTS

We are grateful to staff at The University of Edinburgh Central Bioresearch Services for assistance with animal care, particularly Michael Walls, Rosie Shiels and Duncan McNeil.

## DATA AVAILABILITY STATEMENT

*Functional data underlying this article can be available on reasonable request to the corresponding author. RNA sequencing data underlying this article have been deposited in the European Nucleotide Archive (ENA) with accession number: PRJEB85116*.

## ABBREVIATIONS

BP: blood pressure.

