## Supplemental Figure Legends for "“Loss of circulating glucocorticoid rhythm disrupts the circadian transcriptome and vascular reactivity in the mouse renal artery.”"

**Supplementary Figure Legends:**

**Supplementary Figure 1: Vasoconstriction response to KPSS (A) or phenylephrine (PE, B) does not exhibit diurnal variation in renal arteries from mice treated with vehicle or corticosterone, but corticosterone treatment reduced the renal artery sensitivity to PE at both ZT0 and ZT12.**

Renal arteries were dissected from male mice treated with vehicle (veh, blank silastic pellets) and corticosterone (cort, corticosterone silastic pellets) and mounted onto an isometric wire myograph at ZT0 (local time 7am) and ZT12 (local time 7pm). The vasoconstriction response to high physiological potassium solution (KPSS, A). The vasoconstriction response to increasing doses of PE was compared during the active phase (ZT12) and inactive phase (ZT0) in renal arteries from (B) vehicle-treated mice and (C) corticosterone-treated mice and was also compared between vehicle and corticosterone-treated mice during the inactive phase (D) and active phase (F). Data are expressed as a % of the maximum KPSS-induced constriction value and are mean ± SD and analysed by matched two-way ANOVA, significant p values are underlined and in italics, n=12

**Supplementary Figure 2: Transcripts retaining rhythmicity under arrhythmic corticosterone, demonstrating changes in acrophase and amplitude.**

Renal arteries were dissected (at 2 hourly intervals over a 48-hour period) from C57BL6 mice treated for 7 days with vehicle or corticosterone (cort) pellets (A) Heatmaps showing the gene expression pattern (shown as Z scores) of rhythmic genes from renal arteries from vehicle treated (left) and corticosterone treated (right) mice with different acrophases or different amplitudes as detected and analysed by CircaCompare. (B) Venn diagram detailing the rhythmic transcripts with changed acrophase (blue), amplitude (red) or both (overlapping). (C) Gene profiles of the major circadian clock genes over 24 hours. White and black bars indicate the inactive (light) and active (dark) phases respectively and each dot represents an individual biological sample (n=2 where each n is a mix of both renal arteries from 2 mice. ZT time is indicated on X axis, were ZT 0 is the time of lights on.

**Supplementary Figure 3: Pearson’s correlation of genes exhibiting de novo rhythms in renal arteries from corticosterone-treated mice.**

Renal arteries were dissected (at 2 hourly intervals over a 48-hour period) from C57BL6 mice treated for 7 days with vehicle or corticosterone (cort) pellets. Transcripts were analysed by LimoRhyde with p value for cosinor regression. Genes with rhythms de novo to the corticosterone group were defined as having a p value for rhythmicity of <0.005 in the corticosterone group and a p value of <0.1 in the vehicle group. Pearson’s correlation heatmap showing the R-values (where R=1.0 is a perfect positive correlation) for correlation between the 20 most rhythmic genes that are exclusively rhythmic in renal arteries from corticosterone-treated mice.
