## Supplementary figures and images for "“Loss of circulating glucocorticoid rhythm disrupts the circadian transcriptome and vascular reactivity in the mouse renal artery.”"

### Supplemental Figure 1

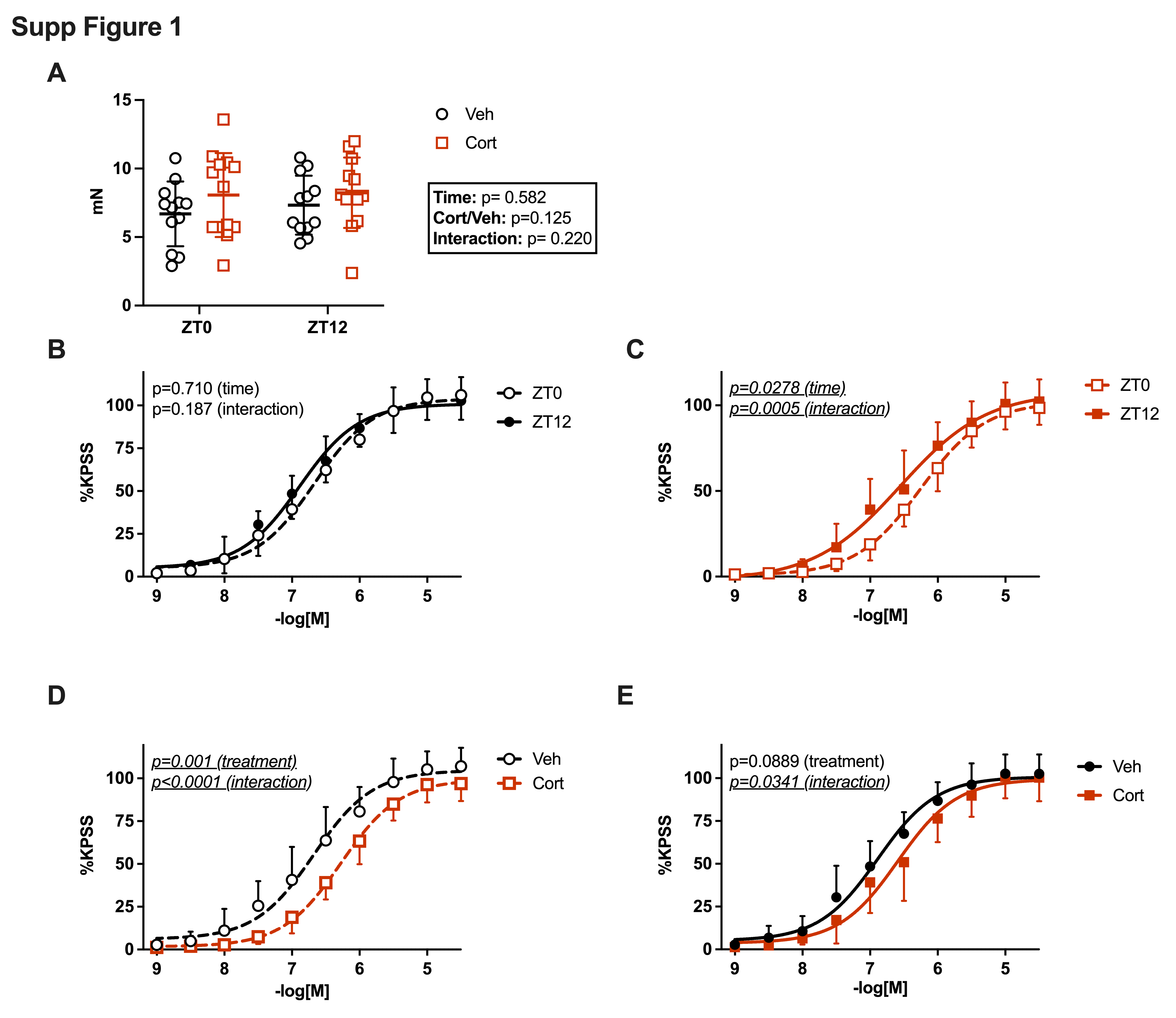

### Supplemental Figure 3

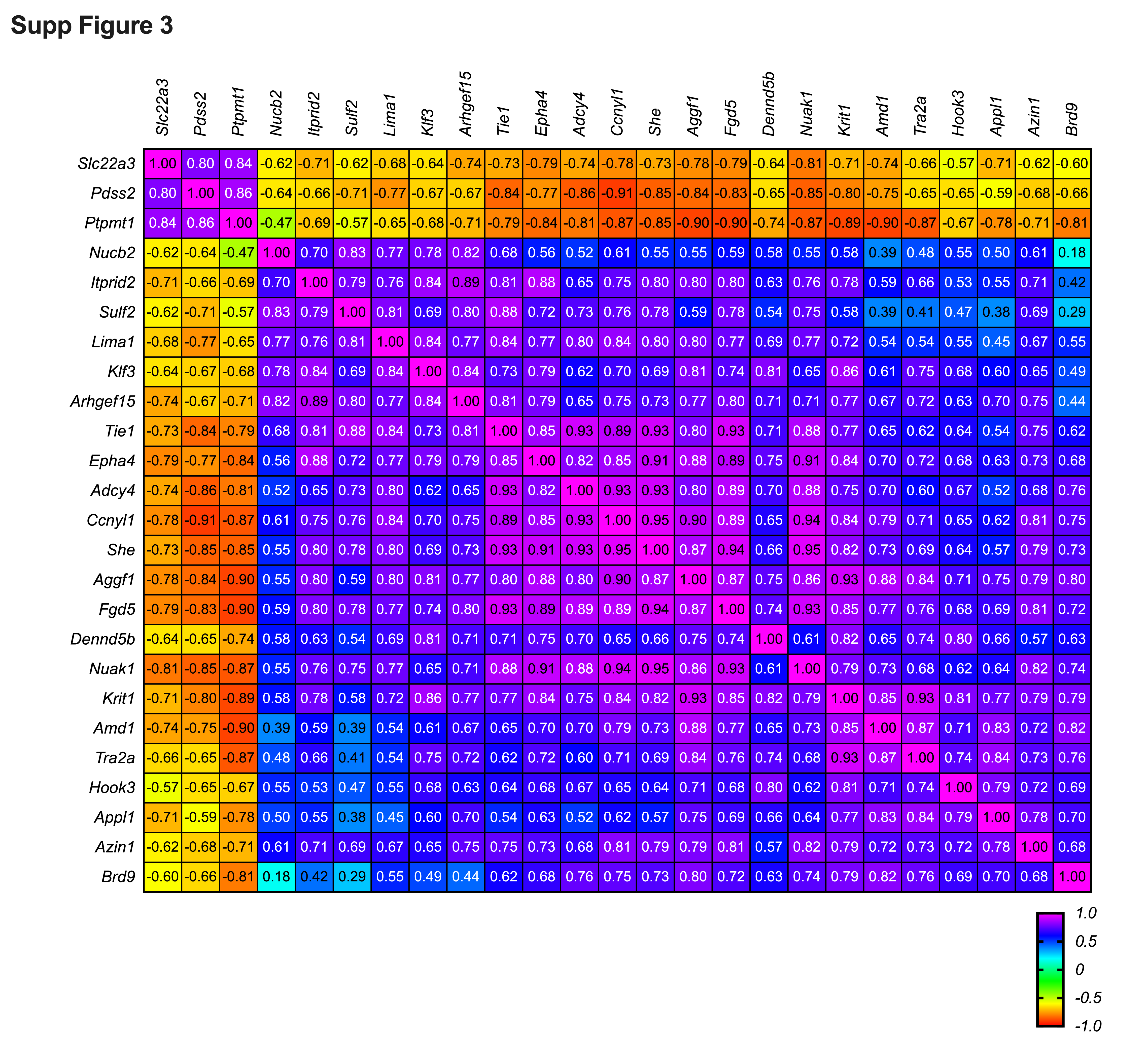
